# Extreme variation in recombination rate and genetic variation along the Sylvioidea neo-sex chromosome

**DOI:** 10.1101/2020.09.25.314054

**Authors:** Suvi Ponnikas, Hanna Sigeman, Max Lundberg, Bengt Hansson

## Abstract

In the majority of bird species, recombination between the sex chromosome pairs in heterogametic females (ZW) is restricted to a small pseudoautosomal region (PAR), whereas recombination is ongoing along the entire Z chromosome in the homogametic males (ZZ). Recombination has a strong impact on the sequence evolution by affecting the extent of linkage, level of genetic diversity and efficacy of selection. Species within the Sylvioidea superfamily are unique among birds in having extended Z and W chromosomes (“neo-sex chromosomes”) formed by a fusion between the ancestral sex chromosomes and a part of chromosome 4A. So far the recombination landscape of the Sylvioidea neo-sex chromosomes remains unknown, despite its importance for understanding sequence evolution. Here, we use linkage mapping in a multi-generation pedigree to assemble, and assess the recombination rate along the entire Z chromosome of one Sylvioidea species, the great reed warbler (*Acrocephalus arundinaceus*). This resulted in an 87.54 Mbp and 90.19 cM large Z including the ancestral-Z, where the small PAR (0.89 Mbp) is located, and the added-Z. A striking result was an extreme variation in recombination rate along the Z in male great reed warblers with high rates at both telomeric ends, but an apparent lack of recombination over a substantial central section, covering 77% of the chromosome. This region showed a drastic loss of nucleotide diversity and accumulation of repeats compared to the highly recombining regions. Nonetheless, the evolutionary rate of genes (measured by dN/dS) did not differ between these regions, suggesting that the efficacy of selection on protein-coding sequences is not reduced by lack of recombination.

## Introduction

Recombination plays a key role in evolution and adaptation, since it creates genetic variation by rearranging alleles into new haplotypes and breaks the linkage between loci allowing them to evolve independently. Therefore, recombination has a critical role in maintaining fitness of large genomic regions, which becomes evident when recombination is lacking (Charlesworth & Charlesworth 2000). In this case, selection has to operate on the level of linked regions instead of independent mutations, which limits the spread of beneficial mutations and the efficiency by which mildly deleterious mutations are being removed, resulting in less effective selection. The population genetic processes linking a lack of recombination and lowered level of adaptation are referred to as Muller’s ratchet and Hill-Robertson interferences (Hill & Robertson 1966, Muller 1964). The former refers to stochastic loss of mutation-free sequence, leading to the accumulation of deleterious mutations. The latter refers to linked selection, where selection for beneficial mutations will fix also linked mildly deleterious mutations (hitchhiking) and purifying selection on deleterious mutations will remove also beneficial mutations (background selection). The end results of these processes, such as loss of genetic variation, accumulation of repeats, change in evolutionary rates and ultimately sequence degeneration, are often observed on heteromorphic sex chromosomes and other evolutionary old non-recombining regions (Charlesworth & Charlesworth 2000).

Besides that recombination has a strong impact on sequence evolution, it can also be under selection itself: depending on the signs and strengths of epistatic fitness interactions between linked genes, recombination can be either selectively beneficial by breaking negative linkage or harmful by breaking co-adapted gene complexes (Butlin 2005). Recombination rates have been observed to vary between species, populations, sexes and individuals (Stapley et al. 2017a). Recombination is also not uniform within the genome, which is reflected by recombination hotspots, where the rate is much higher compared to the general level. These regions can be both stable and dynamic, depending on the taxa and molecular mechanism regulating the recombination events (Stapley et al. 2017b, Sardell & Kirkpatrick 2020). The rate of recombination also depends on the physical position, with centromeric parts often showing lower rates compared to chromosome ends. This bias of recombination events towards telomeres occurs especially in males in vertebrates, while females show more evenly distributed patterns (Sardell & Kirkpatrick 2020).

Heteromorphic sex chromosomes stand out from the general recombination patterns at the genome level. Their evolution is dictated by the lack of recombination between large sections of the sex chromosome pair in the heterogametic sex (XY males and ZW females, in male and female heterogametic systems, respectively), while recombination proceeds in the homogametic sex (XX females and ZZ males). This phenomenon is observed across a wide range of species including animals, plants and fungi (Bachtrog et al. 2014). Even though recombination suppression is often expanding along the sex chromosomes over time, they also tend to retain recombining region(s), known as the pseudoautosomal region (PAR). The persistence of PARs could be due to selection for maintaining recombination or because these regions are essential for the proper segregation of sex chromosomes during meiosis which involves chiasma formation (Otto et al. 2011).

Birds have a female heterogametic sex determination system (ZW females and ZZ males), where females in the vast majority of species do not recombine for most parts of their sex chromosome pair and consequently exhibit a small PAR. There are however exceptions, such as in some palaeognath species (e.g. the common ostrich, *Struthio camelus*), which retain recombination over a substantial part of the sex chromosomes and thus have a large PAR. Also, while the avian sex chromosomes have a deep ancestral homology dating back approx. 100 Myrs, a few lineages have neo-sex chromosomes formed by fusions between the ancestral sex chromosome pair and (parts of) one or more autosomes (Pala et al. 2012b, Gan et al. 2019, Sigeman et al. 2019, Sigeman et al. 2020a). One of these neo-sex chromosome pairs occurs in the songbird superfamily Sylvioidea and was formed approximately 25 Myrs ago when a part of autosome 4A (0-9.6 Mbp) fused to the ancestral sex chromosomes (Pala et al. 2012a, Pala et al. 2012b, Sigeman et al. 2020b). The great reed warbler (*Acrocephalus arundinaceus*) belongs to the Sylvioidea superfamily and earlier studies have presented an incomplete linkage map for the species based on microsatellites and AFLP, including also the Z chromosome (Hansson et al. 2005; Åkesson et al. 2007). Although, the fine-scale recombination pattern across the great reed warbler genome remains largely unknown due to the low marker-density of these linkage maps, the earlier maps strongly suggested that males recombine less than females on the autosomes and that both sexes have markedly shorter map distances compared to other bird species, such as chicken (*Gallus gallus*, Groenen et al. 2009) and collared flycatcher (*Ficedula albicollis*, Kawakami et al. 2014). For example, Dawson et al. (2007) observed linkage map distances of male and female great reed warblers to be only 6.3% and 13.3% of those of the chicken, respectively.

The earlier finding of relatively low level of recombination in the great reed warbler, especially in the homogametic males, together with the fact that the sex chromosome pair does not recombine outside the small PAR in the heterogametic females, suggest very low recombination rates along the species’ Z chromosome. This may generate relatively large evolutionary units of co-segregating loci for selection and drift to act upon, which would in turn affect the efficiency of selection and amount of genetic diversity in the great reed warbler sex chromosomes. Thus, studying the rate and pattern of recombination in relation to sequence polymorphism can help to evaluate the causes and consequences of recombination for sequence and genome evolution.

A draft genome assembly is available for the great reed warbler in which 22 scaffolds have been assigned to the Z chromosome using male-to-female coverage ratio and heterozygosity differences of re-sequenced individuals (Sigeman et al. 2020b). However, the physical order of the scaffolds is still lacking. Here we use linkage mapping in a multigenerational pedigree to assemble the entire Z chromosome of this species. We describe the recombination landscape along this chromosome, and evaluate how the recombination rate varies depending on the chromosome position (central vs. telomeric parts). Then, we analyse the association between recombination rate and genetic diversity, linkage disequilibrium and evolutionary rate (measured by dN/dS). The autosome–sex chromosome fusion forming the neo-Z chromosome (and neo-W chromosome) in our study species provides an opportunity to study the temporal aspect of the causes and consequences of recombination, since the fused part of autosomal origin has been sex-linked for a much shorter evolutionary time compared to the ancestral Z (referred as added- and ancestral-Z, respectively, from now on).

## Materials and methods

### Pedigree and DNA samples

We used multi-generation pedigree data from a long-term study population of great reed warblers at Lake Kvismaren in Sweden. The study species’ ecology, as well as monitoring and sampling practices, are described in e.g. Bensch et al. (1998), Hansson et al. (2005; 2018) and Åkesson et al. (2007). The pedigree data was collected between the years 1984 and 2004 (see Hansson et al. 2018), and for the present analyses we divided the data into 79 three-generation families (grandparents, parents and offspring). As some birds were present multiple times in the pedigree (e.g. as either parent or offspring), the full number of birds in data was 511, of which we had genotype data from 221 individuals. DNA was extracted from field-collected blood samples using a standard phenol/chloroform protocol, and quantified with Nanodrop.

### Genomic data

Single digest (Sbf1) restriction site-associated DNA (RAD) library constructing, sequencing (Illumina 100 bp paired-end), trimming and demultiplexing was performed by BGI (Hong Kong; see Hansson et al. 2018). The sequence reads were mapped to the female great reed warbler reference genome (Sigeman et al. 2020b) with the mem algorithm (Li 2013) in bwa v. 0.7.15 (with default settings except for the –M flag to mark shorter split reads as secondary to ensure downstream compatibility with picardtools). The output was converted into a bam format and sorted with samtools 1.4. Duplicate reads were removed with picardtools v. 2.6.0 (http://broadinstitute.github.io/picard) and the remaining reads were realigned with GATK 3.8-0 (McKenna et al. 2010).

To find high-confidence genetic markers, we called and filtered single nucleotide polymorphisms (SNPs) with three different pipelines (Supplementary Material 2) and then included only those SNPs in the final data set that were found in all three filtered outputs. As erroneous SNPs can cause map inflation and wrong marker ordering, we wanted to aim for as error free data as possible. After filtering, we had 50 265 markers left in 3005 scaffolds (covering 1 200 165 392 bp). Ten of these scaffolds were known to be Z-linked based on their sex-specific coverage and synteny to zebra finch Z chromosome (scaffolds 2, 5, 31, 134, 94, 98, 4139, Sigeman et al. 2020b).

We checked the Mendelian inheritance consistency in the pedigree individuals with VIPER (Paterson et al. 2012) using a sample of 10 000 loci, which showed less than 6 % genotype error in the majority of individuals, with the exception of ten birds with higher genotype errors (6-18%) that suggested incorrect pedigree relationships. We checked the parentage for these ten birds in Cervus v. 3.0.7 (Kalinowski et al. 2007), which corrected the pedigree position for five of them. The remaining five birds with uncertain parentage were removed from the data. Thus, the final data consisted of 240 individuals (146 offspring).

### Linkage map for Z chromosome

A linkage map was built with Lep-MAP3 (Rastas 2017), which is a software for constructing ultra-dense linkage maps with genome-wide pedigree data. The ParentCall2 module in Lep-MAP3 identified a high proportion of Z-linked markers (being homozygous in females) on seven scaffolds (scaffolds 2, 5, 31, 134, 94, 98, 4139) that already had been identified as Z-linked through analyses of resequenced female and male genomes (Sigeman et al. 2020b). This previous analysis categorized scaffolds as Z-linked if they had either (i) a female-to-male genome coverage of <0.55 or (ii) a female-to-male genome coverage between 0.55 and 0.65 and an absolute difference between female and male heterozygosity scores >0.1 (Sigeman et al. 2020b). Moreover, our Lep-MAP3 analysis identified one additional Z-linked scaffold that the previous analysis failed to detect (scaffold 92, size 203298 bp).

To localize the border between the sex-linked part of Z and the PAR, we calculated sex-specific heterozygosity and coverage for all Z-linked SNPs in vcftools v. 0.1.14 using the pedigree birds (Supplementary Figure 1). Eight SNPs showed heterozygosity and coverage patterns in line with a situation where a W-copy had been mapped to the Z (both F/M coverage and female heterozygosity close to 1). We excluded these SNPs from further Lep-MAP3 steps, as they could interfere with the linkage mapping. We also removed markers within sex-linked Z-scaffolds (identified with 0.5 F/M coverage ratio) that were not recognized as sex-linked by Lep-MAP3. This was a conservative approach as we expect all sex-linked SNPs to be homozygous in females due to the loss or divergence of the W-copy. The remaining number of markers was 50257 overall and 850 on the eight Z-scaffolds.

We used the module SeparateChromosomes2 with LOD score 16 to form linkage groups (LGs). The sex-linked markers were initially assigned to two LGs, but with lower LOD score these SNPS assigned into one group. Thus, the division into two linkage groups was concluded to be artificial and we combined all Z-markers into one LG for the further steps.

Additional markers were assigned to LGs in the JoinSingles2 module, by lowering the LOD score from 15 to 3 while simultaneously controlling the quality of linkage with lodDiff-setting, which prevents the markers from assigning to more than one LG with equally high likelihoods. The last step of assigning new markers (LOD=3 and lodDiff=2) was repeated twice. We ran ten independent marker orderings (starting from random order each time) in the module OrderMarkers2 and the one with the highest likelihood value was chosen. Here, we used grandparent genotypes to phase the data. We used Kosambi mapping function and set recombination to occur only in males (recombination2=0). This latter setting applies to most of the Z, but, it is not optimal for the PAR region where both sexes recombine. To be certain that this setting did not interfere with the marker ordering and the following anchoring of the scaffolds (see below), we also ran the marker ordering and anchoring without the assumed PAR scaffold (Supplementary Material 1).

Since the recombination landscape varies between sex-linked Z and PAR (only males recombine in the former and both sexes in the latter), we also ran the marker ordering step separately for the PAR (9 SNPs) to be able to use more suitable settings (i.e., allow both sexes to recombine) for this region. However, the OrderMarkers2 was unable to find the best marker order for the PAR as all ten replicate runs had identical likelihood values, but different marker orders. Therefore we cannot present exact recombination rates for both sexes in this region. In this case, we therefore used marker order according to the physical map order along the scaffold.

### Anchoring the Z scaffolds

The scaffold ordering and orientation was done in ALLMAPS (Tang et al. 2015) with default settings based on the male linkage map (best marker order) for the Z chromosome. The software added a 100 bp gap between each of the scaffolds.

### Recombination rate

To get the most accurate estimates on recombination rates, we re-evaluated the genetic distances for the final physical order of markers (N=664) based on their anchored positions in the ALLMAPS output. Genetic distances were re-evaluated in OrderMarkers2 module but the order itself was not evaluated anymore. Markers without anchored position in the final Z chromosome were excluded at this point. The local recombination rate along the chromosome was estimated in *MareyMap* web-service (at http://lbbe-shiny.univ-lyon1.fr/MareyMapOnline/; Siberchicot et al. 2015, Siberchicot et al. 2017), where we fitted LOESS (span 0.2) to describe the relationship between the genetic and physical distances.

### Recombination in relation to genome characteristics

We extracted the chromosomal region within the first and last markers in the male-specific Z linkage map and divided this region into non-overlapping 200 kbp bins (N=427). The PAR region was also divided into 200 kbp bins (N=4), but since we did not get accurate map distances for this interval, PAR is included only in the comparisons between the recombination regions (see below). Bins overlapping with scaffold gaps were not included in the analyses.

We used a fitted LOESS to interpolate genetic distance point estimates for the start and end position of each bin within the male-specific Z (i.e., excluding PAR). The bin-specific recombination estimate was then the genetic distance (cM) within these point estimates. All recombination estimates within the non-recombining region of the Z chromosome (between 8.47 and 76.42 Mbp, see below) were set to zero.

We studied recombination in relation to several variables expected to be affected thereof: genetic diversity (measured as the nucleotide diversity; pi), linkage disequilibrium (r2), Tajima’s D, proportion of exonic and repeat sequences and GC content. Genome characteristics were calculated for the 200 kbp bin data.

Linkage disequilibrium, genetic diversity (pi) and Tajima’s D were estimated from five whole-genome resequenced great reed warbler males (150 bp paired-end Illumina Hiseq X, Sigeman et al. 2020b) from the same population as the pedigree samples. For details regarding extraction, library preparation and trimming of raw reads, see Sigeman et al. (2020b). Mapping of trimmed reads to the reference genome and subsequent removal of duplicated reads followed the same pipeline as used for the RAD sequencing libraries (see above). Variants were called on the Z chromosome using freebayes with default settings. The raw set of variants was filtered using a combination of vcftools (Danecek et al. 2011) and vcflib (Garrison 2012). The filtering steps included removal of variants overlapping annotated repeats or that had a mean coverage more than twice the median mean coverage of all variants. We further required that alternative alleles should be supported by at least one read on each strand (SAF>0 & SAR>0) and at least one read centered to the left and the right (RPL>0 & RPL>0), and that the quality of the variant should be more than 20. Genotypes with less than 10x coverage were recorded as missing and sites with more than 20 % missing data were filtered. Finally, complex variants and multi-nucleotide SNPs (MNPs) were decomposed into separate SNPs, and indels and bi-allelic SNPs were selected for downstream analyses. Bcftools (Li 2011) was used to extract variants located in each window for which recombination rate had been estimated. For each window, vcftools was used to calculate Tajima’s D and nucleotide diversity. To obtain measurements of linkage disequilibrium (r2), we used beagle version 5.1 (Browning and Browning 2007) to phase and impute missing genotypes. For the phased data we used vcftools to calculate pairwise r2 between SNPs that were at least 1 kb from each other within each window and only retained comparisons with r2 ≥ 0.1. For each window we calculated a mean r2 across all pairwise SNP comparisons.

Other genome variables (GC content, exon proportion and repeat proportion) were calculated from the annotated reference genome (Sigeman et al. 2020b). Exon and repeat proportions were calculated as the proportion of the bin that was covered by each sequence type in bedtools v. 2.26.0 (overlapping features were collapsed).

By utilizing the results of the recombination landscape, and sex-specific coverage and diversity (from both re-sequencing and RAD-seq data), we defined the border between PAR and sex-linked Z, and further divided the chromosome into different recombination regions (Fig. 1A-B and Supplementary Figure 1). Relationships between recombination and genome characteristics were first studied by comparing the three recombination regions of Z: PAR (where both sexes recombine), male-recombining (MREC) and non-recombining (NONREC; see Fig. 2 and Results), which should describe the effect of lacking recombination particularly. The relative amount of recombination between these regions is assumed to be: PAR > MREC > NONREC. The effect of the amount of recombination was then tested by correlations within the MREC region, which was done also separately for the ancestral- and added-Z. The physical position effect on recombination was tested by correlation with the distance to chromosome end and by comparing chromosomal regions (ancestral-versus added-Z).

### Gene-specific evolutionary rates

In order to assess the efficacy of selection in relation to recombination, we calculated gene-specific evolutionary rates (dN, dS and dN/dS ratio) between great reed warbler Z-linked protein-coding genes (N=383) and their zebra finch orthologues as described in Sigeman et al. (2020b). Here we included also the PAR genes (N=16) in the analyses. The dN/dS ratio compares the amount of non-synonymous substitutions at non-synonymous sites to synonymous substitutions at synonymous sites, and tests if a gene is under positive selection (dN/dS > 1), purifying selection (dN/dS < 1) or evolve neutrally (dN/dS = 1). The dN/dS rate is often observed to correlate negatively with recombination rate, which results in higher dN/dS values in regions of low recombination, as purifying selection can not remove slightly deleterious mutations effectively due to Hill-Robertson interference. The recombination rate estimate for each gene’s midpoint was inferred from the fitted LOESS (using a span of 0.1). Relationships between recombination and evolutionary rates were first studied by comparing the three recombination regions of Z and then by correlating the amount of recombination with the evolutionary rates within sex-linked Z.

## Results

### Linkage map for Z

The Z chromosome linkage group (LG-Z) included 676 markers located on ten scaffolds. All eighth scaffolds identified as Z-linked in ParentCall2 module (scaffolds 2, 5, 31, 94, 92, 98, 134 and 4139) were assigned to LG-Z. In addition, two scaffolds with autosomal patterns regarding female/male read coverage and SNP heterozygosity (scaffolds 181 and 217), as would be expected for markers in the PAR, were assigned. Indeed, scaffold 217 (with nine assigned markers) represents the PAR in the great reed warbler Z chromosome (Fig. 3) as it, in addition to the pattern of linkage, has a coverage and diversity similar to autosome, shows synteny to the zebra finch scaffold ‘Z random’ (Z-linked but not ordered scaffolds) and the collared flycatcher’s PAR contigs (Smeds et al. 2014). Scaffold 181, however, was assigned to LG-Z with only one marker (corresponding only to 0.1% of its total marker number) and showed synteny to chromosome 1A in zebra finch, great tit (*Parus major*) and collared flycatcher, and it was therefore not included in LG-Z. Alignment of the great reed warbler PAR and the first sex-linked scaffold (scaffold 92) with the flycatcher PAR and the first 1.2 Mb of the zebra finch Z chromosome (Rhie et al. 2020) is shown in Fig. 3A.

Moreover, scaffold 98 showed a chimeric pattern indicative of a mis-assembly. One part had markers assigned to LG-Z and showed synteny to the fused part of chromosome 4A (i.e. added-Z), while markers on the other part were left unassigned and showed synteny to chromosome 19, 6 and 1 random in the zebra finch. This scaffold was therefore split into one Z-linked part (scaffold 98a) and one autosomal part (scaffold 98b), of which only 98a was included in LG-Z (this also match the data in Sigeman et al. (2020b). A small region (110 kbp) in the beginning of scaffold 98a still shows coverage data corresponding to autosomal patterns (Fig. 1A), but none of the autosomally behaving SNPs within this region got assigned to Z either. This region showed synteny to chromosome 1 random in zebra finch, so instead of representing a part of PAR this scaffold still seems to have a small misassembly.

### Anchoring the Z scaffolds

ALLMAPS arranged the Z scaffolds in the following order and orientation: 98a(+) 31(-) 5(+) 2(-) 134(-) 92(-) 217(+). The order remained the same when the PAR scaffold (217) was excluded (see Supplementary Material 1). The anchored scaffolds correspond to 87 535 819 bp of the Z chromosome. Scaffolds 94 and 4139 were assigned to the Z linkage group, but ALLMAPS could not anchor them, as they both had only 1 SNP. These scaffolds show sex-linked coverage (F/M coverage of 0.5) and synteny to Z chromosome in zebra finch, great tit and collared flycatcher, so we classified them as “Z random” (known to belong to Z, but not placed on the final map). Based on their coverage, synteny information and lack of variation, their likely positions are in the NONREC region. For the figures and the results below, we present the Z chromosome in an order where the ancestral-Z is at the beginning and the added-Z in the end of the chromosome, i.e. 217(-) 92(+) 134(+) 2(+) 5(-) 31(+) 98a(-), as this follows the direction of previous avian Z chromosomes (Fig. 1B).

**Figure 1.**
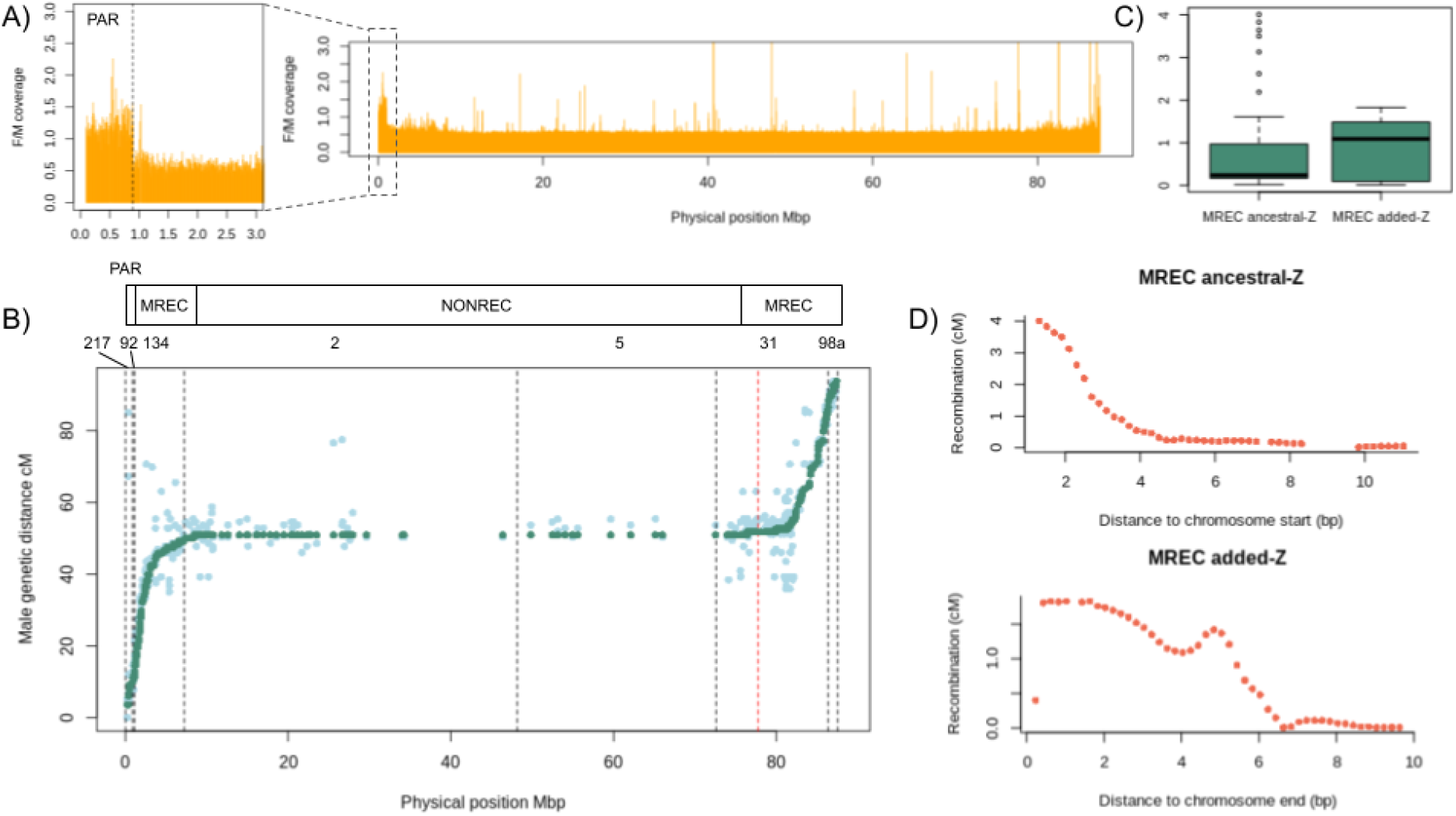
The anchored (ordered and oriented) great reed warbler Z chromosome and its recombination landscape. A) Female-to-male coverage ratio (based on 10 re-sequenced individuals) along the Z chromosome showing the border between the sex-linked region and the PAR (latter zoomed in). B) Male genetic distance as a function of the physical distance. Genetic marker order for the full Z chromosome, which was used to order the Z-linked scaffolds, is shown in light green. Re-evaluated genetic distances based on the final physical positions of markers are shown in dark green (evaluated only for the anchored Z scaffolds). All genetic map positions are based on male recombination. Ancestral part of the Z is in the beginning of the chromosome (starting with PAR) and the added region is in the end, and the red dashed line shows the approximate border between the ancestral- and added-Z (sizes 77.9 and 9.6 Mbp, respectively). Black dashed lines mark divide scaffolds (217, 92, 134, 2, 5, 31 and 98a). Above the plot is a schematic illustration of the three recombination regions of great reed warbler Z chromosome: PAR (where both sexes recombine), male-recombining regions at the both ends (where only males recombine, MREC) and non-recombining region (no recombination in neither sex, NONREC). C) Recombination rate within MREC compared between ancestral- and added-Z. D) Relationship between recombination rate and distance to chromosome ends within MREC, presented separately for the ancestral- and added-Z.

### Recombination rate

The re-evaluation of genetic distances for the full Z gave a map length of 90.19 cM, but the settings were not fully suitable for PAR in this estimate. The sex-linked Z (excluding PAR) got a total map length of 83.69 cM corresponding to a physical region of 86 477 970 bp. Dividing the genetic map length with the physical length gives a general recombination rate of 0.97 cM/Mb across the male-specific Z. However, male recombination ceased completely between 8 473 772 and 76 416 949 bp (at genetic map position 50.90 cM, Fig. 1B). Thus, a 67 943 177 bp region (i.e., 77.6 % of the chromosome) was not recombining, in neither of the sexes, based on our pedigree data. This forms three regions to the recombination landscape of the great reed warbler Z chromosome: PAR (where both sexes recombine), MREC (male-recombining Z) and NONREC (non-recombining Z; Fig. 1B). The average recombination rate was 4.52 cM/Mb, when estimated for the MREC region only.

The MREC region is located in each end of the neo-Z chromosome and can therefore be divided into an ancestral- and an added-Z part. The recombination rate was 5.41 cM/Mb for the former and 3.90 cM/Mb for the latter (i.e., 1.4 times higher in the ancestral-Z). However, when the rates were compared between these regions using the 200 kbp data, they were not statistically different (Mann-Whitney U-test, p-value=0.663, Fig. 1C).

The recombination rate depended strongly on physical position as it increased significantly towards the chromosome ends (Spearman: r^S^= −0.933, p<2.2e-16) and the correlation remained significant also when tested separately for the ancestral- and added-Z MREC parts (r^S^= −0.986, p<2.2e-16, and r^S^= −0.916, p<2.2e-16, respectively). The relationship between recombination rate and chromosomal position showed an almost exponential shape in the ancestral-Z, while in the added-Z pattern was less clear (Fig. 1D).

**Figure 2.**
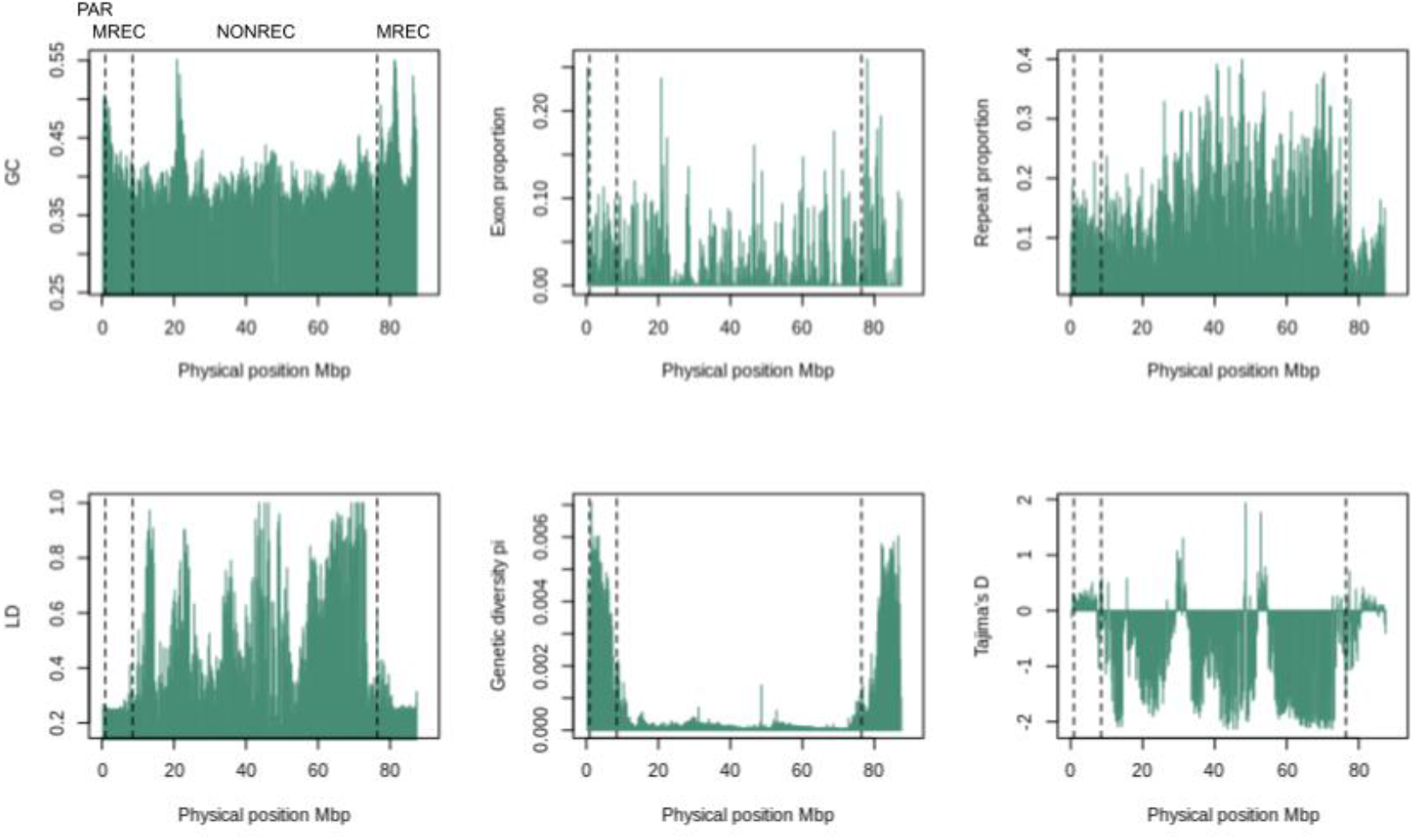
Genome characteristics, i.e. GC content, exon proportion, repeat proportion, linkage disequilibrium, genetic diversity and Tajima’s D, measured in 200kbp bins in relation to the physical position along the great reed warbler Z chromosome. Dashed lines mark the boundaries between the three types of recombination regions: PAR (pseudoautosomal region, where both sexes recombine), MREC (male-recombining region) and NONREC (non-recombining region).

### Recombination rate in relation to genome characteristics

The relationships between genome characteristics (Fig. 2) and recombination were first contrasted between the three recombination regions (PAR, MREC, NONREC; Fig. 4). The GC content was significantly different between all three regions, showing the highest value in the PAR and the lowest in NONREC region. Exon proportion was significantly higher in MREC compared to NONREC. The repeat proportion in turn was significantly higher in the NONREC than in MREC. Linkage disequilibrium was significantly lower in MREC and PAR when compared to the NONREC. Genetic diversity and Tajima’s D were in turn significantly higher in MREC and PAR compared to the NONREC. Since the PAR had only four 200 kbp bins, its results and comparisons need to be interpreted cautiously due to the small number of data points.

When the genome characteristics were compared between the ancestral- and added-Z part of MREC, only Tajima’s D (Mann-Whitney U-test, p-value=0.0003) and repeat proportion (Mann-Whitney U-test, p-value=2.832e-10) showed a significant difference between these regions (both were higher in the ancestral-Z). The remaining parameters (GC, exon proportion, genetic diversity and linkage disequilibrium) did not differ between the ancestral- and the added-Z parts of MREC (data not shown).

Next, we evaluated the relationship between recombination rate and genome characteristics within MREC. Genetic diversity (Kendall’s *tau* 0.6299, p<2.2e-16) and repeat proportion (Kendall’s *tau* 0.2014, p=0.0054) correlated positively, while exon proportion (Kendall’s *tau* −0.2523, p=0.0005) and linkage disequilibrium (Kendall’s *tau* −0.3278, p=6.1e-06) correlated negatively, with the recombination rate. The remaining correlations were non-significant. When these associations were tested separately for the ancestral- and added-Z (Fig. 5), linkage disequilibrium and genetic diversity retained a strong negative and a strong positive correlations, respectively, in both regions (Fig. 5D-E). However, the negative correlation between exon proportion and recombination was found only in the added-Z part of MREC (Fig. 5B). Both regions had positive association between proportion of repeats and recombination rate (Fig. 5C).

**Figure 3.**
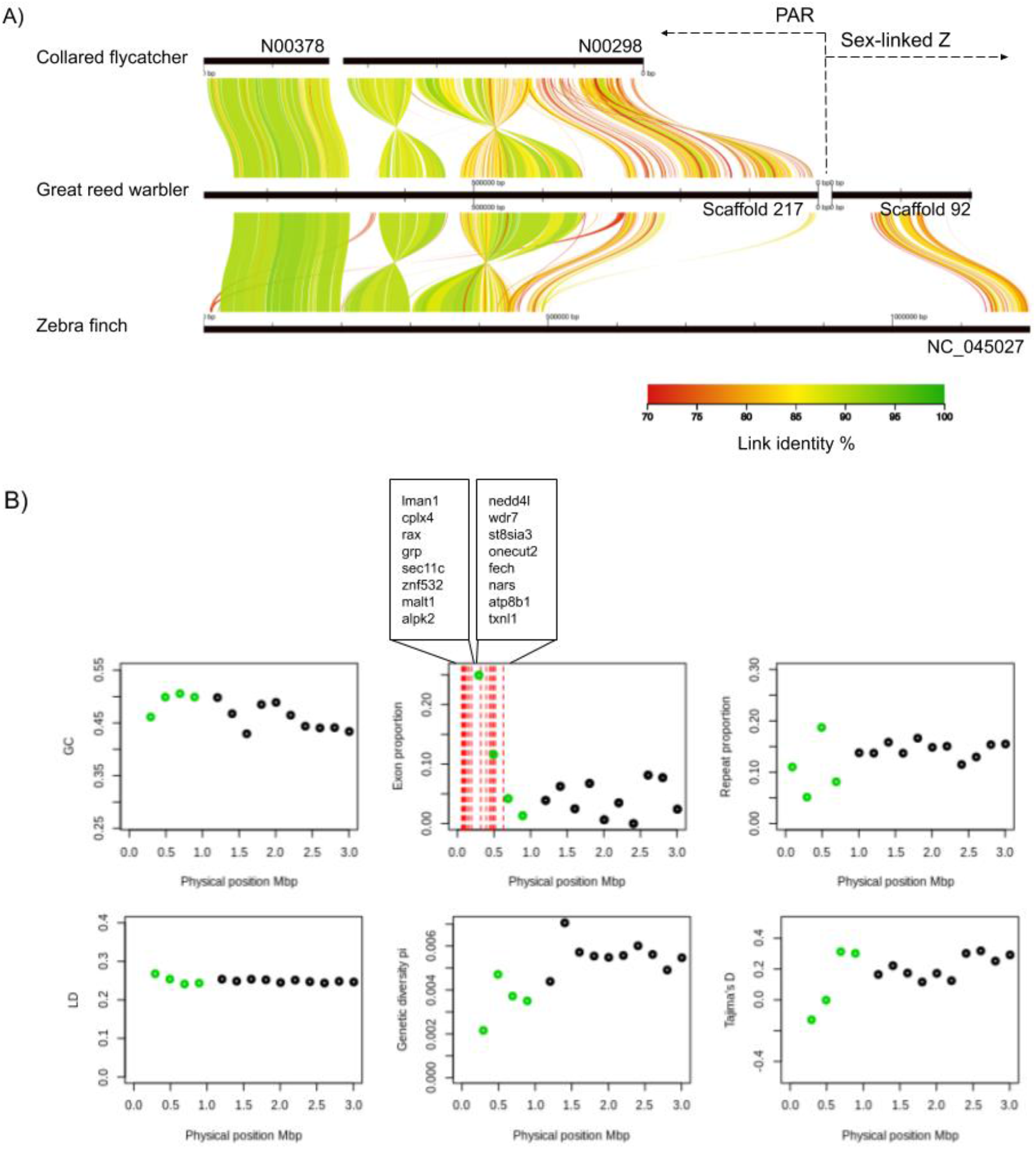
The great reed warbler PAR (scaffold 217) and the beginning of the sex-linked region (scaffold 92) A) aligned to the previously identified flycatcher PAR scaffolds N00298 and N00378 (N02597 is not shown as it did not show synteny to our PAR) and the first 1.2 Mb of the zebra finch Z chromosome from the latest genome assembly (scaffold NC_045027 from genome version bTaeGut2.pat.W.v2). The zebra finch scaffold has a long gap (consisting of N’s) between the PAR and sex-linked regions. The alignment was plotted using the AliTV v.1.0.6 browser tool (Ankenbrand et al. 2017). B) Genome characteristics, i.e. GC content, exon proportion, repeat proportion, linkage disequilibrium, genetic diversity and Tajima’s D, measured in the 200 kbp bins along the great reed warbler PAR (marked with green). For comparison, shown is also the first 10 bins in the sex-linked Z (marked with black). The graph describing the exon proportions shows also the positions for the 16 protein-coding genes in the PAR, which have been listed corresponding to their physical order in the sequence.

Due to the somewhat unexpected observation of a positive correlation between recombination rate and repeats, we further analysed major types of repeats separately: DNA transposons, LINEs, SINEs, LTRs, unknown repeats, low complexity repeats and simple repeats (Supplementary Figure 2). In the comparison between the three regions, LINEs and LTRs were significantly higher in NONREC compared to both PAR and MREC, whereas simple repeats were significantly higher in NONREC than in MREC (Supplementary Figure 2B). When comparing ancestral- and added-Z within MREC, only LINEs were significantly different between the two regions (Mann-Whitney U-test, p-value=3.567e-14, higher in ancestral-Z). Correlations with recombination rate within MREC were significant only for simple repeats (Kendall’s *tau* 0.3060, p-value=2.417e-05). This result was observed also when tested separately for the ancestral- and added-Z (Supplementary Figure 2C). The rest of the significant correlations showed differences between the regions: LTRs were positively correlated in the added-Z and negatively in the ancestral-Z. Unknown repeats had a positive correlation with recombination rate only in the added-Z and LINEs only in the ancestral-Z (Supplementary Figure 2C).

**Figure 4.**
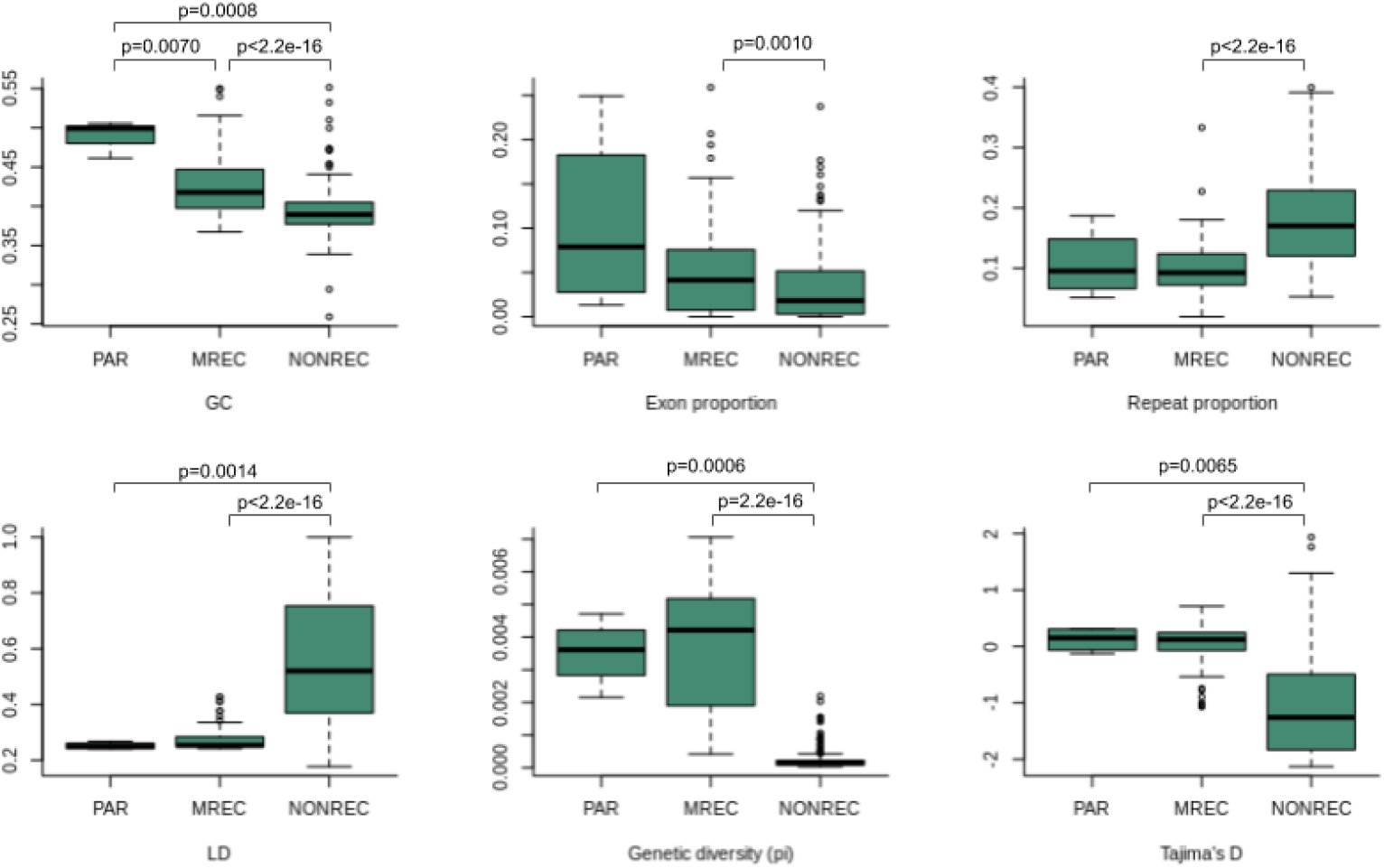
Comparison of genomic variables (GC content, exon proportion, repeat proportion, linkage disequilibrium, genetic diversity and Tajima’s D) between the three recombination regions (PAR, MREC and NONREC) of the great reed warbler Z chromosome. Definitions for the recombination regions are: PAR (pseudoautosomal region, where both sexes recombine), MREC (male-recombining region) and NONREC (non-recombining region). All data is evaluated in 200 kb bins. P-values are shown for the statistically significant differences (Mann-Whitney U-test, p-value < 0.05) between regions.

**Figure 5.**
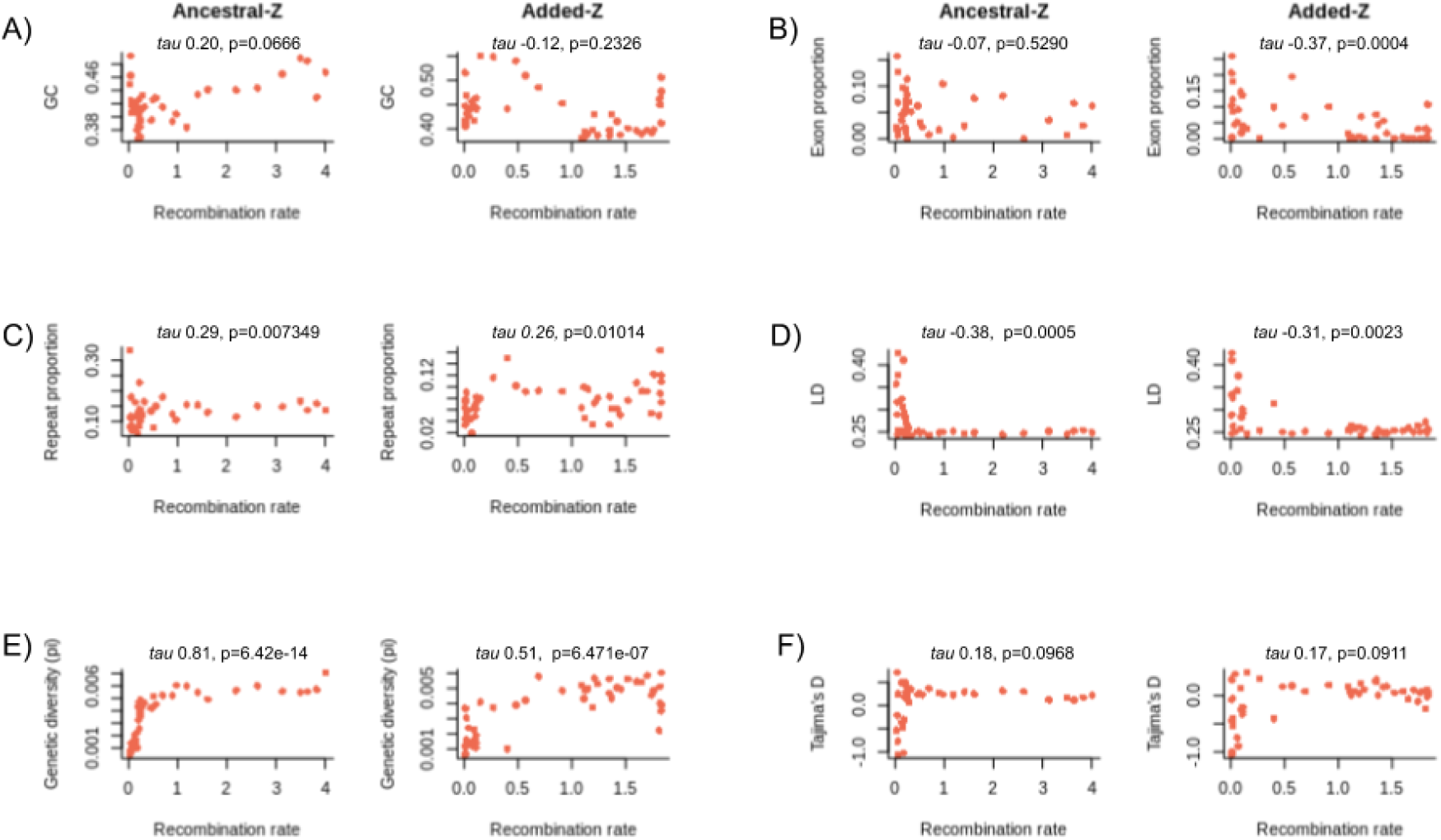
Correlations between recombination rate (cM/200kbp) and genomic variables (GC content, exon proportion, repeat proportion, linkage disequilibrium, genetic diversity and Tajima’s D) on the great reed warbler Z chromosome, assessed in 200kb bins, within both ancestral- and added-Z. Correlations were tested with Kendall’s *tau*.

### Gene-specific evolutionary rates

Whether a Z chromosome gene has lost its gametologous W-copy or not is strongly associated with its evolutionary rates (dN, dS and dN/dS, Sigeman et al. 2020b), and we took this into account by analysing genes with and without a W-copy, separately. For genes with a remaining W-copy, none of the rates (dN, dS and dN/dS) differed significantly between MREC, NONREC and PAR. Rate comparisons for genes without W-copy showed significantly higher dS (Mann-Whitney U-test, p-value=0.0004897) in MREC compared to NONREC. Next we ran the same comparisons, but separating MREC into ancestral- and added-Z. For genes with W-copy, both dN and dN/dS were significantly higher in PAR compared to the MREC ancestral-Z (Mann-Whitney U-test, p-value=0.045 and p-value=0.032, respectively). For genes without a W-copy, dN/dS was significantly higher in MREC added-Z compared to both NONREC and MREC ancestral-Z (Mann-Whitney U-test, p-value=0.03916 and p-value=0.03078, respectively). MREC added-Z showed also significantly higher dN compared to NONREC (Mann-Whitney U-test, p-value=0.0262). Lastly, dS was significantly higher in MREC ancestral-Z than in NONREC (Mann-Whitney U-test, p-value=0.0001148). When we tested the gene-specific correlations between the amount of recombination and evolutionary rates (within sex-linked Z), none of the correlation were significant for genes with W-copy, whereas in genes without W-copy the rate correlated positively with dS (Kendall’s *tau*=0.1854, p-value=0.00015).

## Discussion

### Recombination landscape in great reed warbler Z chromosome

Based on our pedigree data, most of the great reed warbler Z chromosome (77.6 %) is not recombining at all in either of the sexes (female-specific W lacks recombination). This creates three distinct regions within the chromosome: non-recombining Z (NONREC), male-recombining Z (MREC), and the small PAR where both sexes recombine. The non-recombining region is probably even larger (up to 79 %) as the two un-anchored sex-linked scaffolds (94 and 4139) most likely originate from this region. In a typical case of heteromorphic sex chromosomes, the chromosome pair does not recombine in the heterogametic sex, but in our study species recombination along a large section of the Z chromosome is also absent in the homogametic sex.

The sampled offspring represent only a proportion of the total amount of meioses and potential crossing over events in the population, so we cannot exclude the possibility of rare recombination events within the NONREC area. However, the drastic loss of diversity and accumulation of repeats within the central parts of the Z chromosome are in line with no or very low amount of recombination. The crossing over events cluster at the chromosome ends, where the recombination rate showed further fine-scale dependence on the chromosomal position by increasing towards the ends. This pattern is in line with the general bias of recombination towards the telomeres in male vertebrates (Sardell & Kirkpatrick 2020). When compared to other bird species, the zebra finch shows a similar strong telomeric effect in its recombination rates and has a comparable recombination desert in the central parts of its Z chromosome (Backström et al. 2010). A less pronounced telomeric effect (without the centromeric recombination deserts) have been observed also in chicken (Groenen et al. 2009) and in the collared flycatcher (Kawakami et al. 2014). It is generally thought that each chromosome arm requires at least one chiasma (resulting as a minimum of 50 cM distance) for efficient chromosome pairing and segregation during meiosis. As our results show a shorter total map distance (87.54 cM including two arms), we still miss some of the recombination events in the Z chromosome, especially as the genetic map for the PAR could not be resolved with the current data. The previous linkage maps for the study species have found Z chromosome map lengths of 45.3 cM (Hansson et al. 2005), 155 cM (Åkesson et al. 2007) and 156.1 cM (Pala et al. 2012b). The last two of these maps were partly based on AFLP markers, which are difficult to score and filter for genotyping errors as they are dominant. Therefore the previously observed longer map lengths (> 100 cM) were likely inflated due to genotyping errors.

### Great reed warbler PAR

The PAR has recently been described from several avian species (Zhou et al. 2014, Smeds et al. 2014, Xu et al. 2019). Of these species, the collared flycatcher (*Ficedula albicollis*) arguably has the best description for a passerine PAR and our study adds another fine-scale described PAR. The great reed warbler PAR covers approximately 1 % of the total Z chromosome and it seems to be slightly larger (892 kbp) compared to collared flycatcher (630 kbp), which however was suspected to be incomplete (Smeds et al. 2014). After carefully checking the support for each of the PAR annotations, we could confirm 16 protein-coding genes, which are the same as found in the PAR of collared flycatcher (Fig. 3B, Smeds et al. 2014). Alignments to collared flycatcher and zebra finch showed two inversions in our study species’ PAR (Fig. 3A).

The fusion between part of chromosome 4A and the sex chromosomes in the great reed warbler and other Sylvioidea species occurred to the sex-linked end of the Z chromosomes and it does not seem to have created a second PAR. A small region (110 kbp) in this added-Z end of the chromosome shows autosomal patterns in SNPs and coverage data (Fig. 1A), but none of the markers within this region were assigned to the LG-Z when tested. This small region is thus likely misassembled or contains repetitive sequence and shows autosomal patterns due to that instead of ongoing recombination between Z and W.

### Recombination rate in relation to genome characteristics

The apparent lack of recombination in the central part of the Z chromosome shapes the evolution of this region drastically compared to the recombining parts, which is seen as higher linkage disequilibrium and amount of repeats, as well as a lower GC content. This region had also a lower proportion of exonic sequence, but concluding that this is due to gene degeneration should be done cautiously as the higher amount of repeats in NONREC could partly decrease the proportion of exons in the bins of a certain size. In addition, centromeric regions (which make up part of NONREC) tend to have lower gene density in general. Perhaps the clearest consequence from the apparent lack of recombination is the loss of genetic diversity. The level of genetic variation is determined by mutations, recombination, genetic drift, selection and demography. The neutral expectation of sex-linked Z chromosome diversity corresponds to 0.75 of the general autosomal level, which reflects the difference in the effective population sizes of these genomic regions (Z chromosome N_E_ is ¾ of the autosomal N_E_, assuming equal sex-ratios). The fact that males have a higher mutation rate compared to females (male mutation bias; Kirkpatrick & Hall 2004) can increase this Z/A neutral expectation above the 0.75, as the Z chromosome spends ⅔ of its evolutionary time in males. On the other hand, higher variance in reproductive success in males can decrease this relative diversity below the 0.75, through the decrease in the males’ effective population size (N_E_-males < N_E_-females). Assuming that the PAR reflects roughly the general autosomal diversity level, the male-specific Z of our study species shows a relative diversity of 0.27 compared to the PAR. If purely neutral, a ratio this low would need an unrealistically low N_E_-males/N_E_-females, too low to be explained by the polygynous mating system and potential for male-biased reproductive variance in great reed warblers. When the relative Z/A diversity was calculated separately for MREC and NONREC regions, the former showed equal amounts of variation compared to autosomes (1.04, which is higher than expected), while the latter had drastically lower value (0.06). The similar level of variation between MREC and PAR (despite having different N_E_s) could result from male-mutation bias in the former. The difference in relative diversity values between NONREC and MREC results most likely from the observed lack of recombination and the following linked selection in the former, since the male mutation bias, effective population size, drift and demography should affect these two regions in similar manner.

The negative Tajima’s D values in the NONREC supports the idea of linked selection (selective sweeps and/or background selection) causing a loss of polymorphism. Tajima’s D compares two genetic diversity estimates, based on average pairwise differences and number of segregating sites, and values below zero follow from excess of low frequency polymorphisms. The sex-linked Z is mostly hemizygous in females due to the gene loss in the W chromosome (within NONREC 87.8 % of genes we used in the evolutionary rate analysis have lost its W-copy), which enhances the effect of selection further, since also recessive alleles are exposed to selection in females. We cannot say whether it is mostly purifying or positive selection (or mixture of them) that causes the linked selection and the resulting negative Tajima’s D values. It is a challenging task in general to distinguish between background selection and hitchhiking in causing linked selection (Stephan 2010).

There were three intervals within NONREC that showed high positive Tajima’s D values and thus stood out from the general pattern of negative values in this region (Fig. 2). Closer assessment of two of the wider intervals suggested that high Tajima’s D values could have been caused by divergent, likely introgressed, haplotypes that are present at high frequency in the study population. Due to the low recombination rate in these regions, one would expect that such introgressed chromosomal chunks could be quite long. The third, rather narrow interval, however, is more likely caused by collapsed paralogous sequences as it showed clearly higher coverage.

Linkage disequilibrium is naturally strongly affected by the crossing over events, which break the linkage associations between loci. Our results were in line with this and the LD was significantly lower in NONREC and had a strong negative correlation with recombination rate. The positive association between recombination and GC content in region comparisons was expected too, since the crossing over events are known to affect the nucleotide composition of the sequence through GC-biased gene conversion (Mugal et al. 2013) and higher GC content is linked to recombination hotspots in many species across eukaryotes (Stapley et al. 2017a). In addition to the significantly lower genetic diversity in NONREC, diversity increased strongly with the amount of recombination within MREC. A positive correlation between genetic diversity and recombination is widely observed (Frankham 2012) and both selection and neutral mechanistic processes can explain this association. In non-recombining regions, like in the great reed warbler NONREC, linked selection explains the loss of variation, whereas in recombining regions the positive association could be additionally explained by the crossing over-associated higher mutation rate (i.e., neutral process). This is thought to result from the fact that repairing of the DNA double-strand breaks during the crossing over event is mutagenic (Kulathinal et al. 2008, Arbeithuber et al. 2015).

The proportion of exons and repeats showed expected patterns in the region comparisons as exon proportion was lower and repeat proportion higher in the NONREC. A positive correlation is often found with gene density and recombination, which can be beneficial through reduction in the Hill-Robertson interference between the loci (Hill & Robertson 1966). However, in contrast, the correlation within added-Z was negative, which could suggest selective advantage on the gene linkages (see below). The higher repeat density in NONREC most likely results from the decreased efficiency of selection to remove these sequences in the absence of recombination. Again, added-Z showed an opposite pattern (positive correlation), which we discuss below. Associations between different repeat types and recombination showed complex results. LTRs had the expected pattern with recombination, as they were accumulated in NONREC and showed negative correlation with recombination rate also in the ancestral-Z. Added-Z though, had an opposite positive correlation between LTRs and the rate, so some other unknown factors seem to be affecting this relationship too. Region comparison supported accumulation of LINEs in the absence of recombination too, but when correlated with the rate within MREC, ancestral-Z showed positive association. Thus, some types of repeats seem to accumulate in the absence of recombination as expected, but that is not the general pattern describing the relationship between recombination and repeats in great reed warbler Z.

### Comparing the ancestral- and added-Z

The crossing over events seemed to happen equally often in ancestral- and added-Z within the MREC. However, the recombination rate correlations with distance to chromosome end had slightly different patterns between these regions (Fig. 1D). The rate within the ancestral-Z shows clear dependence on the physical position, which could suggest that the rate variation is not selectively driven by the gene content. The pattern along the added-Z in turn shows some deviation from the exponential association between physical position and recombination rate. This could be caused by many factors. For example, a lack of informative SNPs in the bins where the rate drops from the expected could be one simple explanation. Selection could be another reason for this deviation, if the fitness effect of gene linkage varies along the added-Z. Then selection could create a local rate that is higher or lower than expected based on the physical position. This possible difference in selective regimes between ancestral- and added-Z was supported by significantly lower Tajima’s D in the added-Z. While the values were mostly positive in the ancestral-Z, half of the added-Z region showed negative values (Fig. 2), suggesting linked selection. Selectively advantageous gene linkages in added-Z are further supported by the significant negative correlation between gene density and recombination rate. Further studies are needed to assess this possible variation in the fitness effect of recombination along the Z chromosome.

In addition to the possible selective differences (due to the gene contents) between ancestral- and added-Z, the age difference in the sex-linkage could affect sequence patterns between these regions. In addition to the Tajima’s D, none of the other genome characteristics differed significantly between these regions. However, some of the unexpected correlation patterns in the added-Z could indeed be caused by the younger sex-linkage of this region. For example, the lack of correlation between recombination and GC content in the added-Z could be explained by age difference. If the current recombination landscape of added-Z differs from the previous autosomal one, the GC content could still reflect both, previous and current recombination rate effects. This could disrupt the sign of ongoing GC biased gene conversion and the following positive correlation between recombination rate. The positive correlation between recombination and repeats in the added-Z could also be explained by this signature of previous and current recombination rate landscape or alternatively it follows from the observation that recombination is occurring outside the gene dense regions (i.e., where repeat proportion is higher) in the added-Z.

### Gene-specific evolutionary rates

Even though the lack of recombination and the following linked selection has left a strong signature on genetic diversity, the efficacy of selection in protein-coding sequence does not seem to have been hampered since dN and dN/dS had similar values between NONREC and MREC. The lack of association was further strengthened by the non-significant correlations between dN, dN/dS and recombination rate. Similar kind of lack of association between recombination and adaptation has been observed for example in primates (Bullaughey et al. 2008). The positive correlation between dS and recombination in genes without W-copy was the only significant association and it could be caused by the mutagenic effect of recombination and/or the GC-biased gene conversion that is associated with recombination. It could have been expected that NONREC shows higher rates as a result of relaxed purifying selection (i.e., there is a negative association between recombination and evolutionary rate), but instead our results suggest that there is no relationship between recombination and evolutionary rates. If there are rare recombination events within the NONREC region that our pedigree data did not record, it could be enough to enable selection to work efficiently, especially as the gene density is lower within the NONREC, which could decrease the effect of linked selection on genes further. Another explanation could be that as our substitutions were not calculated separately for great reed warbler (i.e., we did not use a phylogenetic branch-specific analysis) they could partly hide the signal between recombination and substitutions occurring specifically in our study species.

For genes with remaining W-copy, the PAR (where all genes are diploid) showed the highest and MREC-ancestral the lowest rates of protein evolution and these differences (in dN/dS and dN) were also significant. This result is similar to the recent finding in Paleognatheous birds, where some of the species showed higher evolutionary rates in the PAR genes compared to the autosomes and sex-linked genes (Xu et al. 2019a). The great reed warbler PAR region could show higher rates due to higher gene density, which was twice as high (0.1051) compared to the MREC-ancestral (0.0508, see also Fig. 3). This could lead to a situation, where the dense gene organisation is hindering selection to work independently on each gene despite the assumed high recombination rate, causing the genes to accumulate more mutations due to linked selection. This explanation is supported by lower Tajima’s D values and slightly higher LD in gene rich PAR bins (Fig. 3). In genes without W-copy the MREC-added showed highest dN and dN/dS values and these were significantly different compared to the rest of the male-specific Z (dN/dS) and to NONREC (dN). This difference could be explained by different selective pressure (eg. positive selection on recessive beneficial mutations in females) on added-Z genes and/or by the fact that the comparison in these genes was between hemizygous genes in added-Z and diploid autosomal (chromosome 4A) genes in zebra finch.

## Conclusions

We used linkage mapping to assemble 87.54 Mbp neo-sex chromosome in one Sylvioidea species, the great reed warbler (*Acrocephalus arundinaceus*), including and describing also the pseudoautosomal region (0.89 Mbp) for the first time for this species. The great reed warbler Z chromosome shows an extreme variation in male recombination rate along its length with high rates at telomeric ends, but an apparent lack of recombination over a substantial central section that covers 77% of the chromosome. This variation has left strong signatures on the sequence evolution, especially to the level of genetic diversity, linkage disequilibrium and GC content. The central non-recombining region showed a drastic loss of nucleotide diversity, driven by linked selection. Nonetheless, the evolutionary rate of genes (measured by dN/dS) was not influenced by the amount of recombination, suggesting that the efficacy of selection on protein-coding sequences is not reduced by lack of recombination.

## Acknowledgements

Samples were collected non-destructively and with the appropriate permissions from the Malmö-Lunds djurförsöksetiska nämnd (M 45-14 and 17277-18). Re-sequencing was performed by the SNP&SEQ Technology Platform in Uppsala, which is part of the NGI Sweden and SciLife-Lab, supported by the Swedish Research Council and the Knut and Alice Wallenberg Foundation. Sequence data (RAD) used in this study are deposited in NCBI Sequence Read Archive under accession PRJNA578893. Bioinformatics analyses were performed on resources provided by SNIC at UPPMAX. The research was funded by grants from the Royal Physiological Society in Lund, the Erik Philip-Sörensen’s Foundation, Stiftelsen Olle Engkvist and the Wenner-Gren Foundations (to S.P.), and the Swedish Research Council (to B.H., consolidator grant no. 621-2016-689).

