## Supplementary Material for "Extreme variation in recombination rate and genetic variation along the Sylvioidea neo-sex chromosome"

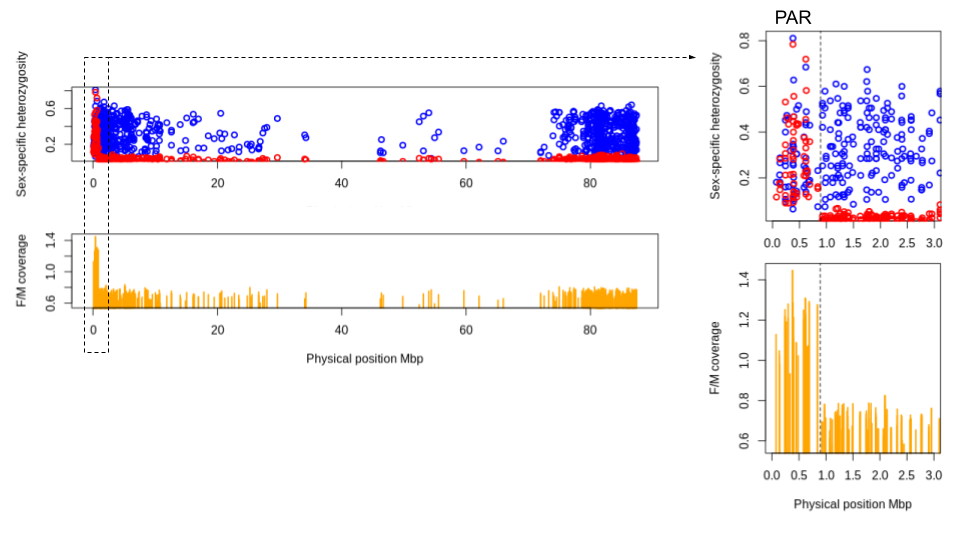
**Supplementary Figure 1**. Sex-specific heterozygosity and coverage based on RAD-seq SNP data along the great reed warbler Z chromosome. Here, the values have been calculated along Z-linked scaffolds that were anchored (ordered and oriented) in ALLMAPS analyses. We have included all filtered markers from these scaffolds (N= 906), i.e., including also those that were not assigned to the Z linkage group, to get best possible coverage across the chromosome. The zoomed in PAR region in the beginning (0-892 kbp, border marked with dashed line) shows an autosomal patterns for both coverage and heterozygosity, while the rest of the chromosome shows clear sex-linked patterns: i) females show half of the mapping depth to the Z compared to males and ii) females are homozygous (since they are hemizygous) for the Z-linked loci. Both of these patterns are caused by divergence or loss of the W-copy in females.

**Current:** 217(-) 92(+) 134(+) 2(+) 5(-) 31(+) 98a(-)

**Without PAR:** 92(+) 134(+) 2(?) 5(+) 31(+) 98a(-)

**Supplementary Material 1**. When the PAR scaffold (217) was excluded, ALLMAPS arranged the Z scaffolds in the following order: 92(+) 134(+) 2(?) 5(+) 31(+) 98a(-). Thus, the order remained the same, but scaffold orientation changed somewhat: the orientation of scaffold 5 flipped and the orientation of scaffold 2 was left unknown. The fact that ALLMAPS cannot orient these two scaffolds with high certainty is expected as they are in the middle of the ancestral Z chromosome, which does not recombine (i.e., genetic map is not informative). When we evaluated the anchored scaffold order by synteny to great tit (*Parus major*) and collared flycatcher, it suggested that the orientation of scaffold 2 (+) is currently most parsimonious, but the orientation of scaffold 5 is more parsimonious in the ALLMAPS output that was ran without PAR scaffold. However, this uncertainty will not affect our recombination rate analyses since scaffold 5 is located in the non-recombining area so its orientation in relation to the neighbouring scaffolds will not change the estimates (see Fig. 1).

**Supplementary Material 2**. Description of the three pipelines used for variant calling and filtering. If not stated otherwise, all filtering steps in different pipelines were performed in VCFtools v. 0.1.14 (Danecek et al. 2011).

In the first pipeline, variant calling was done with freebayes v1.1.0 (Garrison & Marth 2012) and the raw variants output contained 2.55 million SNPs in 3123 scaffolds. Filtering was performed following the SNP filtering approach in dDocent v. 2.2.20 (Puritz et al. 2014), when applicable to our data. We did not include the HWE-filtering step (as our data consists of relatives) or the rad_haplotyper script. Last, we applied two additional filtering steps, which kept only bi-allelic sites and removed sites overlapping with annotated repeats. Repetitive regions were identified by (i) using a bed-file of repeats constructed using a repeat library (fAlb15_rm3.0_aves_hc.lib; provided by A. Suh, Uppsala University) containing Repbase repeats (Bao et al. 2015) from chicken (Hillier et al. 2004) and zebra finch (*Taeniopygia guttata*; Warren et al. 2010), curated hooded crow (*Corvus corone*) repeats (Vijay et al. 2016), and curated collared flycatcher repeats (Suh et al. 2018), and (ii) *de novo* prediction by RepeatModeler (raw output, no manual curation). After filtering, 66 392 SNPs in 344 scaffolds remained. Four individuals were removed after this filtering step because of too much missing data (> 85 %), which left 263 birds in the data set.

In the second pipeline, SNPs were called with freebayes as above, but filtering was different. First we removed any indels, which was followed by removal of sites with quality value below 30, removal of genotypes with less than 8x coverage and with more than twice the average autosomal coverage. After that we removed SNPs with average minDP<15 and average maxDP>200 and sites with more than 50% of missing data. Last, we removed sites with minimum allele frequency below 0.05, SNPs overlapping repeats and kept only biallelic sites. This pipeline resulted in 137 507 SNPs in 622 scaffolds.

In the third pipeline, SNP calling was done with mpileup in samtools v. 1.4 (settings ‐t DP and ‐t SP to keep per‐sample read depth and strand bias, and flags ‐Ato keep anomalous read pairs in variant calling and -g to compute genotype likelihoods and output them in the binary call format) and the call command in bcftools v. 1.6 (with flags -vm to use multiallelic calling model and to output variant sites only). The raw output before filtering consisted of 3.97 million variants. We filtered the data mainly following Hansson et al. (2018), with two additional steps including removal of sites overlapping repeats (see above) and with quality score lower than 30. This led to a remaining 209 126 SNPs in 712 scaffolds.

The final set of SNPs was extracted from the output of the first pipeline (called with freebayes and filtered with dDocent) using bcftools isec by selecting SNPs that were shared by all three pipeline outputs (position and exact allele match required). This data set had 50 614 SNPs in 328 scaffolds. As the female-specific W chromosome cannot be ordered using linkage mapping (as it does not recombine), we removed known W-linked contigs (see Sigeman et al. 2020b) from the data. As the data had been mapped to an earlier version of the reference genome (Sigeman et al. 2020b), we removed all SNPs located on 4884 scaffolds identified as redundant in the final assembly.

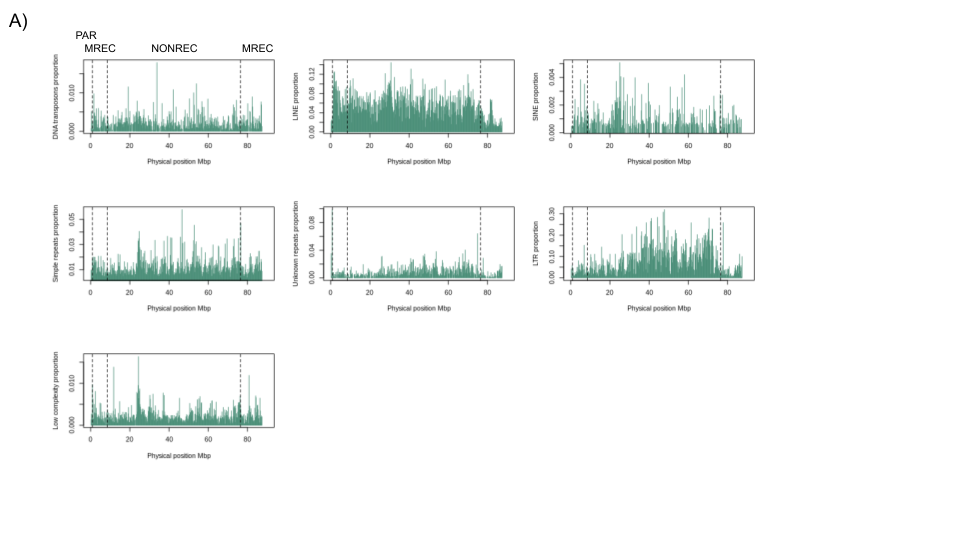

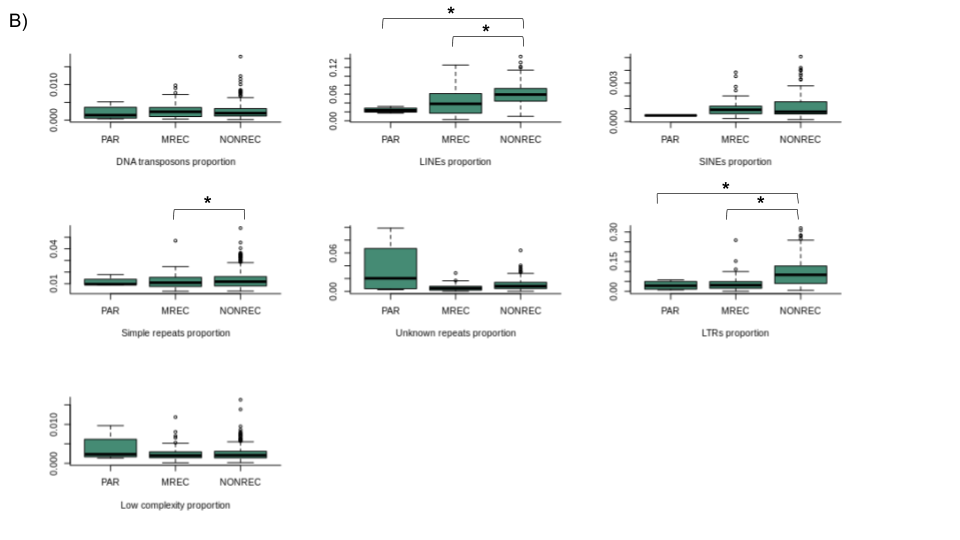

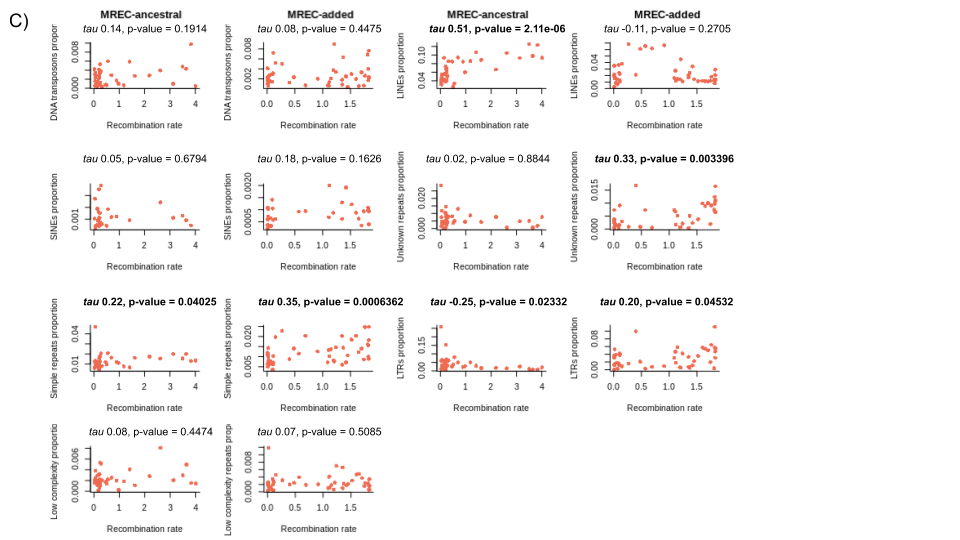

**Supplementary Figure 2**. The repeat variable (proportion of repeats in 200 kbp bins) was divided into different major types (DNA transposons, LINEs, SINEs, LTRs, unknown repeats, low complexity repeats and simple repeats) to further study the relationship between them and the recombination rate. A) Repeat types in relation to the physical position along the great reed warbler Z chromosome. Dashed lines mark the boundaries between the three types of recombination regions: PAR (pseudoautosomal region, where both sexes recombine), MREC (male-recombining region) and NONREC (non-recombining region). B) Comparison of repeat types between the three recombination regions (PAR, MREC and NONREC). Statistically significant differences (Mann-Whitney U-test, p-value < 0.05) between regions are marked with asterisks. C) Correlations between recombination rate (cM/200 kbp) and repeat types, evaluated within ancestral- and added-Z. Correlations were tested with Kendall's *tau* and statistically significant values are highlighted in bold.
